# The sRNA MicC downregulates *hilD* translation to control the SPI1 T3SS in *Salmonella enterica* serovar Typhimurium

**DOI:** 10.1101/2021.08.03.454913

**Authors:** Fatih Cakar, Yekaterina A. Golubeva, Carin K. Vanderpool, James M. Slauch

**Author notes:** Corresponding Author., Department of Microbiology, University of Illinois, B102 Chemical and Life Sciences Laboratory, 601 S Goodwin Avenue, Urbana Illinois, 61801.

## Abstract

*Salmonella enterica* serovar *Typhimurium* invades the intestinal epithelium and induces inflammatory diarrhea using the *Salmonella* pathogenicity island 1 (SPI1) type III secretion system (T3SS). Expression of the SPI1 T3SS is controlled by three AraC-like regulators, HilD, HilC and RtsA, which form a feed-forward regulatory loop that leads to activation of *hilA*, encoding the main transcriptional regulator of the T3SS structural genes. This complex system is affected by numerous regulatory proteins and environmental signals, many of which act at the level of *hilD* mRNA translation or HilD protein function. Here, we show that the sRNA MicC blocks translation of the *hilD* mRNA by base pairing near the ribosome binding site. This binding blocks translation but does not induce degradation of the *hilD* message. Our data indicate that *micC* is transcriptionally activated by SlyA, and SlyA feeds into the SPI1 regulatory network solely through MicC. Transcription of *micC* is negatively regulated by the OmpR/EnvZ two-component system, but this regulation is dependent on SlyA. OmpR/EnvZ control SPI1 expression partially through MicC, but also affect expression through other mechanisms. MicC-mediated regulation plays a role during infection, as evidenced by an increase in *Salmonella* fitness in the intestine in the *micC* deletion mutant that is dependent on the SPI1 T3SS. These results further elucidate the complex regulatory network controlling SPI1 expression and add to the list of sRNAs that control this primary virulence factor.

**IMPORTANCE:** The *Salmonella* SPI1 T3SS is the primary virulence factor required for causing intestinal disease and initiating systemic infection. The system is regulated in response to a large variety of environmental and physiological factors such that the T3SS is expressed at only the appropriate time and place in the host during infection. Here we show how the sRNA MicC affects expression of the system. This work adds to our detailed mechanistic studies aimed at a complete understanding of the regulatory circuit.

## INTRODUCTION

*Salmonella enterica* serovar Typhimurium causes inflammatory diarrhea and potentially life-threatening systemic infection, with an estimated global burden of ∼95 million cases per year world-wide (1). Upon oral ingestion, *Salmonella* transits through the stomach to reach the distal ileum of the small intestine, the initial site of colonization (2, 3). *Salmonella* invades the intestinal epithelium of the host and induces inflammatory diarrhea using the *Salmonella* pathogenicity island 1 (SPI1) type III secretion system (T3SS), a needle-like structure that injects effector proteins into the host cell cytosol (4).

The SPI1 T3SS is controlled by three AraC-like transcriptional regulators, HilD, HilC and RtsA, which constitute a complex feed-forward regulatory loop, each activating transcription of the *hilD, hilC*, and *rtsA* genes, as well as activating *hilA*, encoding the transcriptional regulator of T3SS structural genes (Fig. 1)(5). This system is controlled in response to numerous regulatory proteins and environmental signals, many of which act at the level of *hilD* mRNA translation or stability, or HilD protein activity (6, 7). This includes regulation by a number of small RNAs (8).

**FIG 1.**
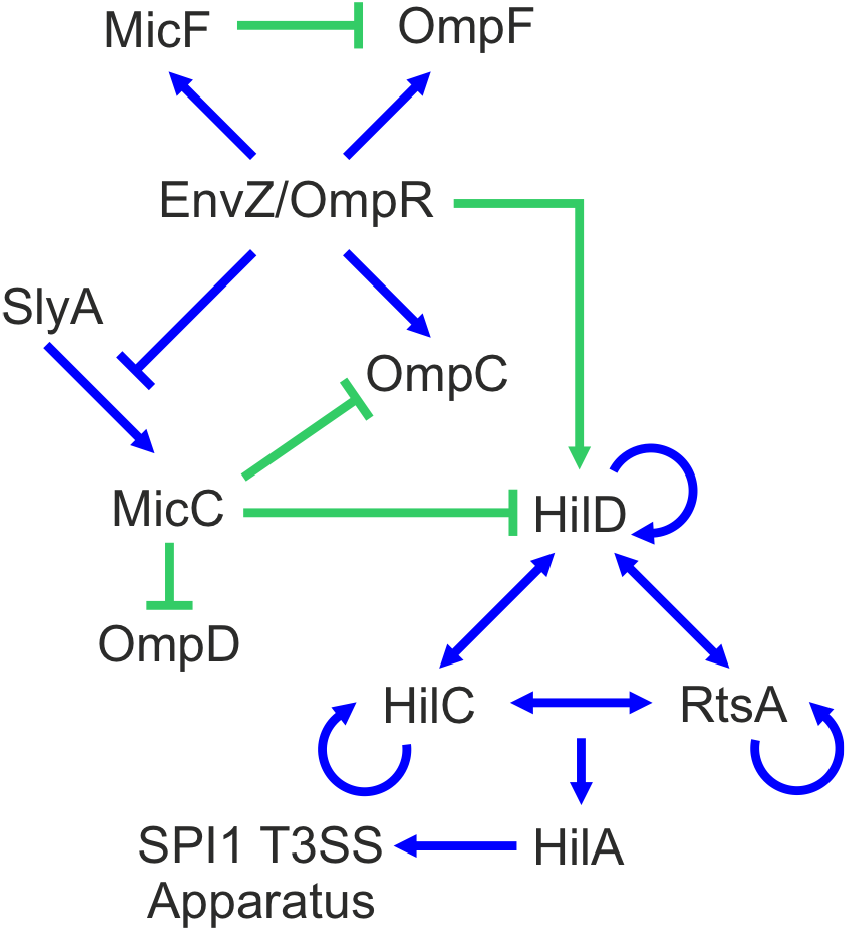
Simplified model of SPI1 T3SS regulatory circuit. Arrows indicate positive regulation, blunt ends indicate negative regulation, blue lines indicate transcriptional regulation, and green lines indicate post-transcriptional regulation.

Small RNAs (sRNAs), generally 50 to 450 bp in length, are increasingly recognized as critical regulators of gene expression (9). Over 300 sRNAs are expressed in *Salmonella*. They play important roles in the regulation of envelope stress responses, metabolism, and virulence. However, the function of most of these sRNAs is unknown or only partially characterized. Many sRNAs control gene expression by imperfect base-pairing with an mRNA near the ribosome binding site (RBS), mediated by the RNA chaperone Hfq (9, 10). This blocks access to the 30S ribosomal subunit, inhibiting translation initiation. In some cases, this also leads to RNaseE-mediated degradation of the message (11).

The OmpR/EnvZ two-component regulatory system was originally characterized as the regulator of the OmpF and OmpC porins in response to changes in osmolarity (12). OmpR/EnvZ is now understood to be a global regulator of virulence in *Salmonella*, activating both the SPI2 and SPI1 type three secretion systems, despite the fact that these systems are primarily required in different niches (13-15). The transmembrane histidine kinase EnvZ autophosphorylates and transfers a phosphoryl group to the response regulator OmpR. At low concentrations of OmpR-P, *ompF* is activated, while at high concentrations of OmpR-P, *ompF* is repressed and *ompC* is activated (16). The sRNA MicF is transcribed upstream and antisense to *ompC* by OmpR. MicF, one of the first sRNAs identified (17, 18), base pairs with the *ompF* mRNA to block translation and facilitate the transition from producing OmpF to OmpC porin in high osmolarity. More recently, the MicC sRNA was identified and characterized as a regulator of the outer membrane porin OmpC in *E. coli* that acts by binding to the *ompC* mRNA near the RBS to prevent 30S ribosome loading (19). In *Salmonella*, MicC downregulates both OmpC and OmpD, binding in the *ompD* coding sequence to initiate RNase-E dependent mRNA degradation (20). Chen et al. (19) reported that *micC* transcription is negatively regulated by OmpR/EnvZ in *E. coli*. Transcriptomic data suggested that *micC* is regulated by OmpR/EnvZ, RpoS, and SlyA in *Salmonella* (21), but regulation of *micC* has not been characterized in detail. SlyA is a member of MarR/SlyA family of bacterial transcriptional regulators. In *Salmonella, slyA* mutants are significantly attenuated in the mouse model of infection (22). SlyA acts both positively and negatively to control expression of some 30 genes (23-26). SlyA controls some genes independently, but often functions in concert with other transcriptional regulators, including PhoP and OmpR (23, 27, 28).

In this study, we define a new regulatory role for MicC sRNA, controlling the SPI1 T3SS in *Salmonella*. We found that MicC base pairs with the leader sequence of *hilD* mRNA to negatively regulate translation of *hilD*. MicC-mediated SPI1 regulation is dependent on environmental signals and regulated through both SlyA and the OmpR/EnvZ two-component system, which acts through or in conjunction with SlyA. We also show that MicC-dependent regulation of SPI1 is important during intestinal infection.

## RESULTS

### The small RNA MicC downregulates the expression of HilD

Several regulatory proteins and signals affecting expression of the SPI1 T3SS act at the level of *hilD* mRNA translation (7). In the few cases that have been characterized, this regulation is mediated either by the RNA binding protein CsrA (29) or sRNAs (30). We previously screened a set of highly conserved sRNAs for those that decrease *hilD* translation when overproduced, and subsequently characterized regulation of HilD translation by FnrS and ArcZ (8). This screen also identified the 109 nucleotide sRNA MicC, first characterized as a regulator of *ompC*, encoding the OmpC porin protein in *E. coli* (19). MicC is encoded in the intergenic region downstream of the pyruvate-flavodoxin oxidoreductase gene (*nifJ*) in both *E. coli* and *Salmonella* and is conserved in the Enterobacteriaceae (Fig. 2A). In *Salmonella*, MicC downregulates translation of the *ompC* and *ompD* mRNAs by base pairing using the highly conserved 5’ 20-30 nucleotides (20).

**FIG 2.**
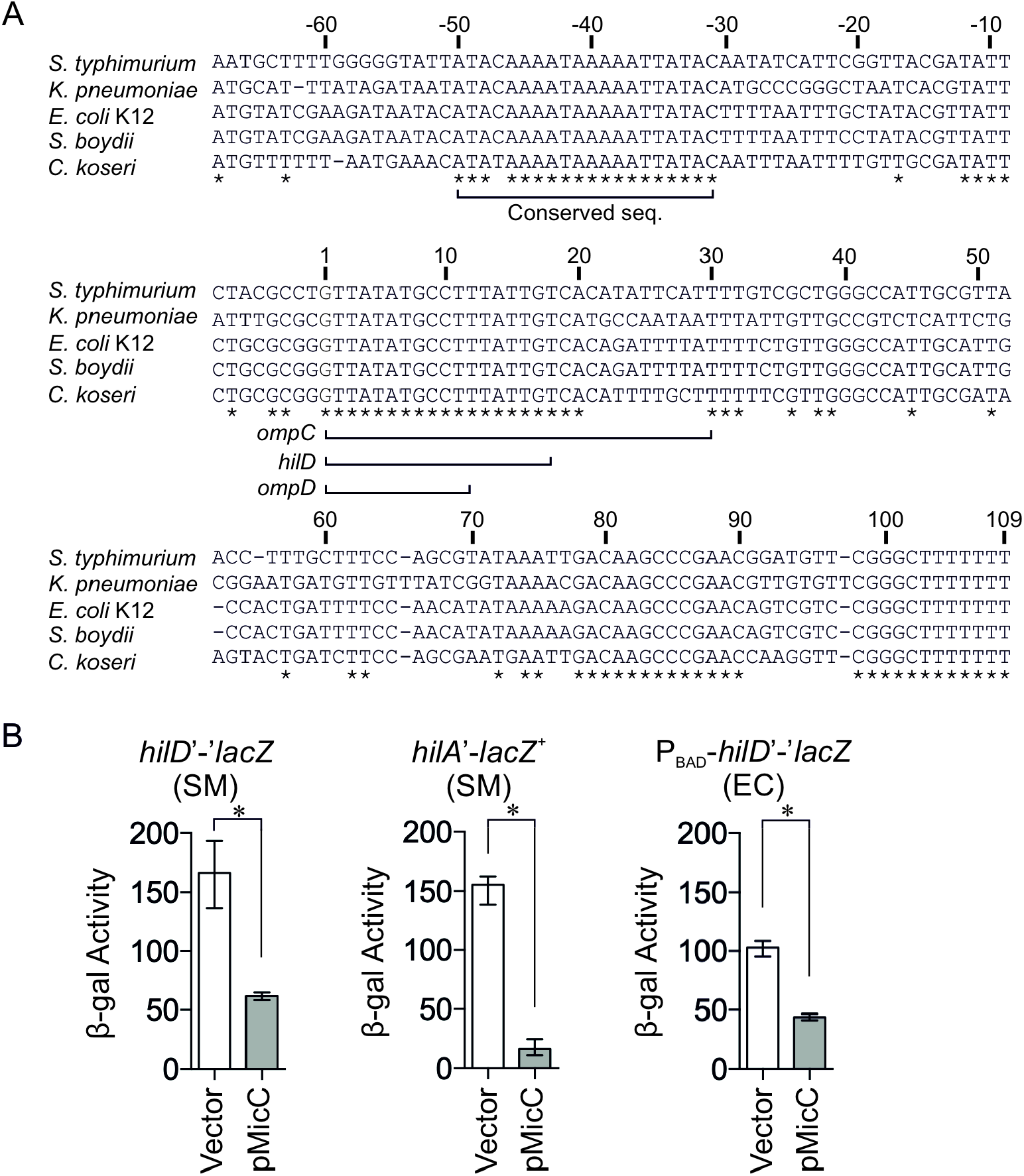
The conserved small RNA MicC negatively regulates the SPI1 T3SS by repressing *hilD* translation in *Salmonella*. (A) Alignment of the MicC sequences from various Enterobacteriaceae. The asterisks indicate sequence identity. Sequences corresponding to the regions of MicC that base pair to *ompC, ompD* and *hilD* are underlined. (B) β-gal activity in *Salmonella* (SM) strains harboring a *hilD*’-’*lacZ* translational fusion or a *hilA*’-*lacZ*^+^ transcriptional fusion, or an *E. coli* (EC) strain harboring a *hilD*’-’*lacZ* translational fusion under control of an arabinose-inducible promoter. Each background contains the pBR-p*lac* vector or pMicC plasmid. β-gal activity units are defined as (µmol of ONP formed min^-1^)×10^6^/(OD_600_ x ml of cell suspension). Results are shown as median with interquartile range and asterisks indicate significant differences between the datasets (n ≥ 4, *P* < 0.05, using a Mann-Whitney test). Strains used: JS749, JS892 and JMS6500, with plasmids pBR-p*lac* vector or pMicC.

To understand the regulation of the SPI1 T3SS system of *Salmonella* by MicC, we overexpressed MicC from the pBRplac plasmid (31) in *Salmonella* strains harboring either an in locus *hilD*’-’*lacZ* translational fusion or a *hilA*’-*lacZ*^+^ transcriptional fusion. Note that the *hilD* fusion strain is a *hilD* null. Thus, this fusion represents the basal level of transcription and is not autoregulated. The *Salmonella* cultures were inoculated in No Salt LB (NSLB) overnight and sub-cultured in High Salt LB (HSLB) for 3 hrs to induce SPI1. The expression of *hilD* was downregulated ∼3-fold in the pMicC strain (Fig. 2B). Expression of *hilA* was decreased 10-fold by MicC (Fig. 2B). These data suggest that MicC negatively regulates HilD expression, leading to a more dramatic effect on *hilA* transcription, consistent with the feed-forward loop model (Fig. 1). To ensure that this is a direct effect on *hilD*, we introduced the MicC plasmid into an *E. coli* strain containing a P_BAD_-*hilD*’-’*lacZ* translational fusion with an arabinose-inducible promoter. The fusion consists of the 35-nt 5′ UTR and the first 11 codons of *hilD* fused in-frame to *lacZ*. Overexpression of MicC in *E. coli* downregulated the expression of *hilD* more than 2-fold (Fig. 2B). These results suggest that MicC acts directly on the *hilD* mRNA to inhibit translation.

### MicC targets the 5′ UTR of *hilD* mRNA by direct base pairing

Bioinformatic analysis using IntraRNA (http://rna.informatik.uni-freiburg.de/IntaRNA/Input.jsp) suggested that MicC has a binding site in the *hilD* mRNA immediately upstream of the ribosome binding site (RBS; Fig. 3A). Based on this prediction, we mutated nucleotides 1 to 5 and 11 to 14 as indicated in the pMicC plasmid. We measured β-galactosidase activity in the *Salmonella hilD*’-’*lacZ* translational fusion strain expressing the MicC mutant (pMicC-mt). The pMicC-mt plasmid conferred no significant repression of *hilD* (Fig. 3B). We then introduced the pMicC-mt plasmid into the *E. coli* P_BAD_-*hilD*’-‘*lacZ* fusion strain. Consistent with the result in *Salmonella*, overexpression of MicC-mt did not regulate *hilD* expression (Fig. 3C). Based on the predicted interaction, we introduced compensatory mutations in nucleotides -18 to -26 of *hilD* in the P_BAD_-*hilD*’-’*lacZ* fusion (Fig. 3A). Overexpression of the wild type MicC had no significant effect on this fusion, whereas the MicC-mt downregulated the mutant *hilD* mRNA (Fig. 3C). These data support the proposed base-pairing interaction between MicC and the *hilD* mRNA. It should be noted that several point mutations and double mutations did not affect the interaction, suggesting that the base-pairing is relatively robust. Similar robust base-pairing interactions were also observed between sRNA MicC and *ompC* mRNA (19) and *ompD* mRNA (20) targets.

**FIG 3.**
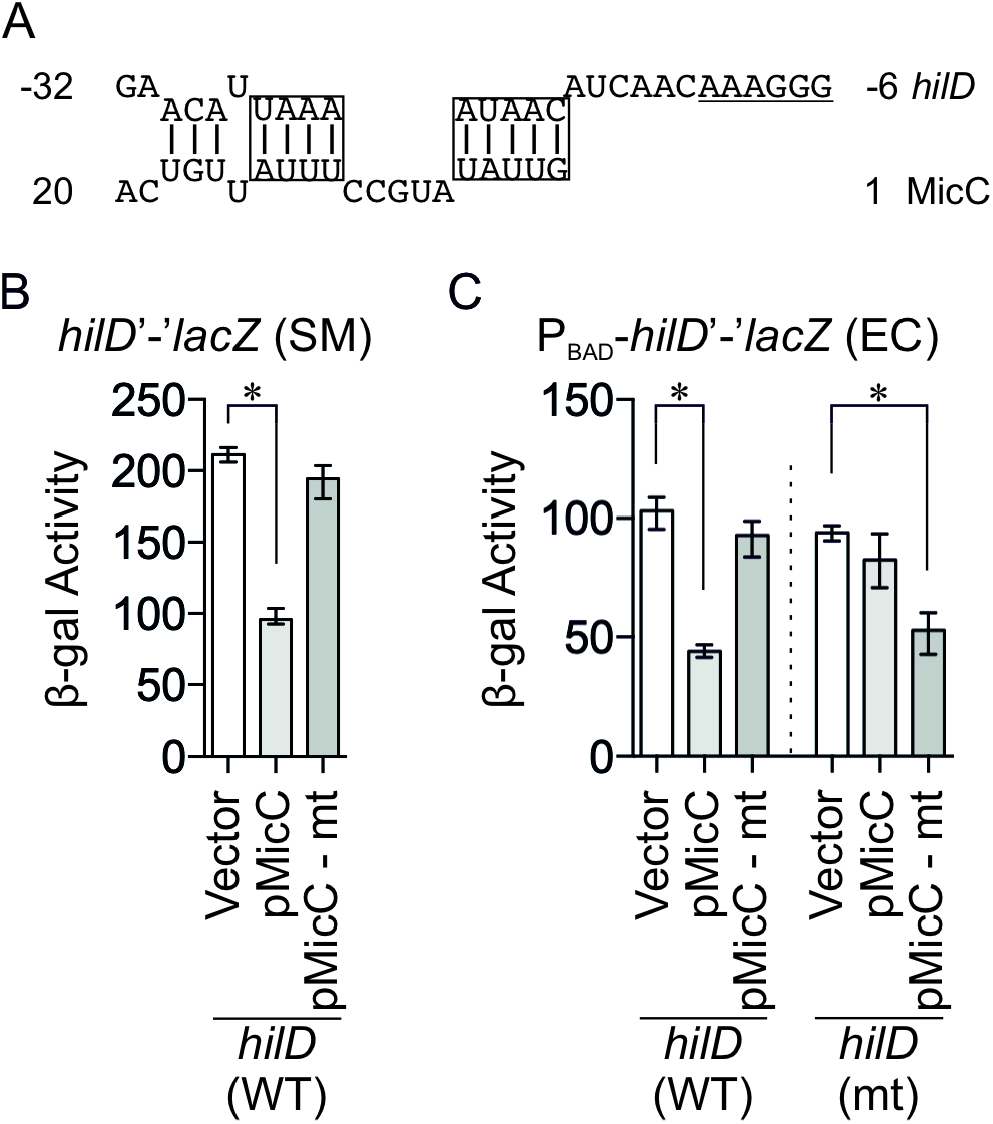
MicC negatively regulates *hilD* translation by base pairing near the RBS of the *hilD* mRNA. (A) Predicted base pairing interaction between MicC and *hilD* mRNA. The RBS is underlined; boxes represent the nucleotides changed in the complementary mutations. (B) β-gal activity in the *Salmonella* (SM) strain harboring the wild type *hilD*’-’*lacZ* translational fusion with vector pBRp*lac*, wild type pMicC, or mutated pMicC-mt plasmid. (C) β-gal activity in *E. coli* (EC) strains harboring either the wild type or mutated *hilD*’-’*lacZ* translational fusion with empty vector, wild type pMicC, or mutated pMicC-mt plasmid. Results are shown as median with interquartile range. Asterisks indicate significant differences between the datasets (n ≥ 4, *P* < 0.05, using a Mann-Whitney test). Strains used: JS892, JMS6500 and JMS6510, with plasmids pBR-p*lac* vector, pMicC, or pMicC-mt.

The data above show that MicC affects translation of *hilD*. We also tested the effect of overexpressed MicC on *hilC* and *rtsA* using translational LacZ fusions in *Salmonella*. In *hilD*^+^ strains, expression of MicC led to a decrease in expression of both *hilC* and *rtsA* (Fig. 4A). However, there was no effect in the *hilD* null background, consistent with the fact that MicC inhibits *hilD* translation (Fig. 4B). Reduced levels of HilD protein decreased transcription of *hilC* and *rtsA*, consistent with the feed-forward loop model (Fig. 1). MicC also did not directly affect translation of either *hilC* or *rtsA in E. coli* (Fig. 4C). MicC downregulates *hilA* transcription via HilD (Fig. 2B). To confirm that MicC does not affect *hilA* translation, we overexpressed MicC in an *E. coli* strain containing a *hilA*’-’*lacZ* translational fusion. MicC had no effect on the expression of this fusion (Fig. 4C). All of these results are consistent with MicC solely regulating *hilD* translation to affect transcription of *hilC, rtsA* and *hilA*.

**FIG 4.**
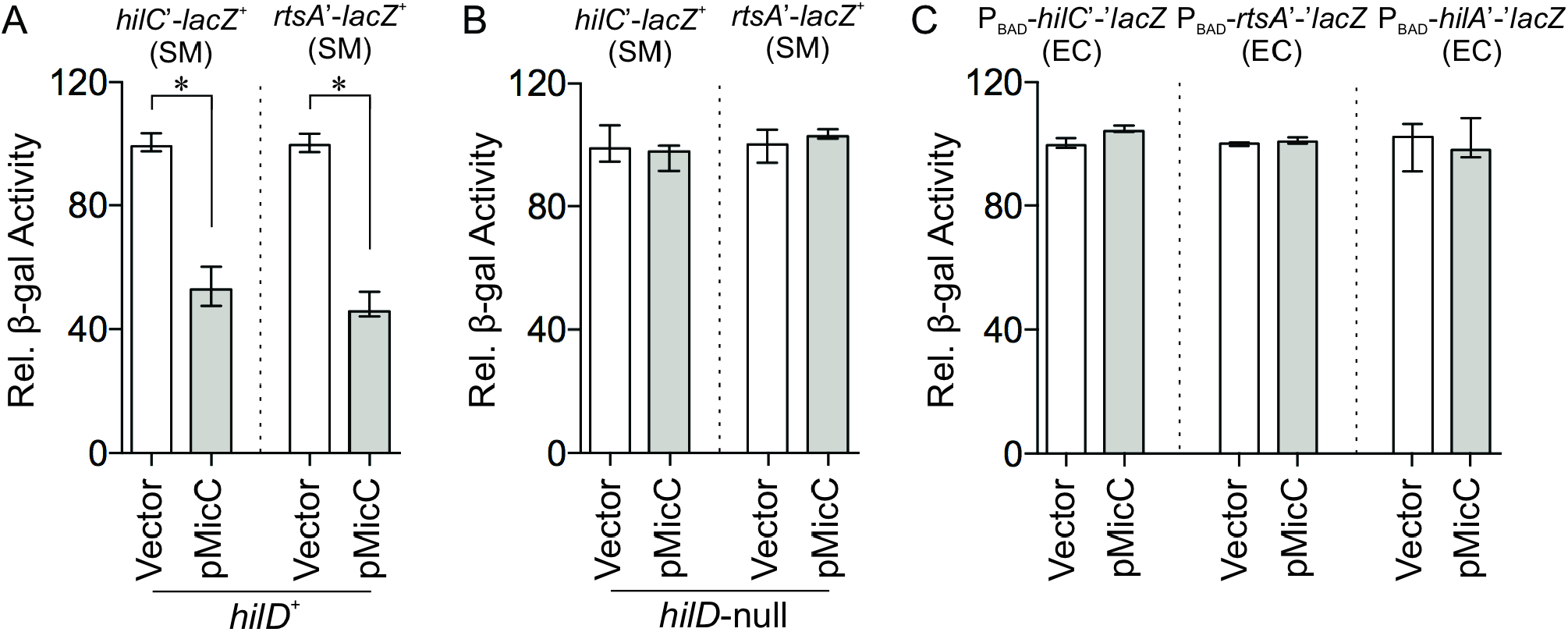
MicC does not regulate HilC, RtsA or HilA. Relative β-gal activity in *Salmonella hilC* or *hilA* transcriptional fusion strains that are (A) *hilD*^+^ or (B) *hilD*-null. (C) Relative β-gal activity in *E. coli hilC*’-‘*lacZ, rtsA*’-’*lacZ* or *hilA*’-’*lacZ* translational fusion strains. All strains include either pBR-p*lac* vector or pMicC plasmid. Results are normalized to each strain containing the vector and are shown as median with interquartile range. Asterisks indicate significant differences between the datasets (n = 4, *P* < 0.05, using a Mann-Whitney test). Strains used: JS2187, JS2196, JS2551, JS2552, JMS6503, JMS6504 and JMS6505, with plasmids pBR-p*lac* vector or pMicC.

### MicC requires Hfq but not RNase E for *hilD* mRNA regulation

The RNA chaperone Hfq is a vital facilitator of sRNA-mRNA imperfect base-pairing (32). To test if the MicC-*hilD* mRNA interaction is dependent on Hfq, we measured *hilD* expression levels in an *hfq* mutant *Salmonella* after introducing the MicC plasmid. There was no significant regulation mediated by MicC in the *hfq* background, suggesting that the interaction and perhaps MicC stability require the Hfq chaperone protein (Fig. 5A). Consistent with our result, Hfq is essential for MicC-dependent regulation OmpC in *E. coli* (19) and OmpD in *Salmonella* (20).

**FIG 5.**
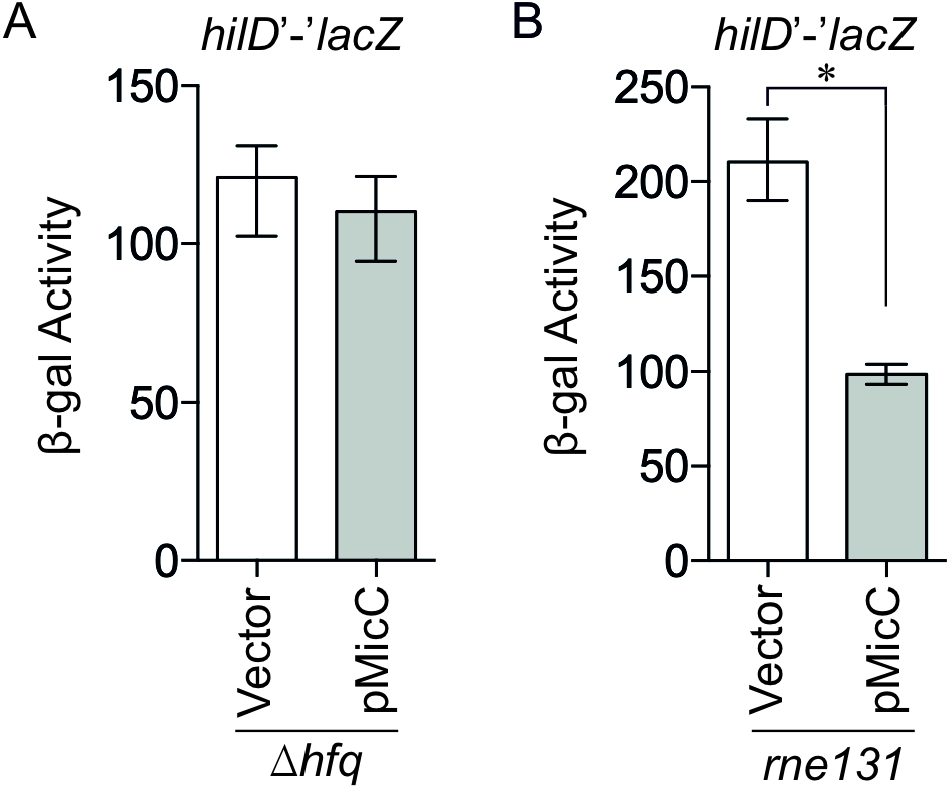
Regulation of *hilD* mRNA by MicC requires Hfq but not RNAse E. β-gal activity in *Salmonella* (A) *hfq* or (B) *rne131* strains harboring the wild type *hilD*’-’*lacZ* translational fusion with vector pBR-p*lac* or wild type pMicC. Results are shown as median with interquartile range and asterisks indicate significant differences between the datasets (n=4, *P* < 0.05, using a Mann-Whitney test). Strains used: JS2118 and JS2119, with plasmids pBR-p*lac* vector or pMicC.

Negative regulation by sRNAs can be due to simple inhibition of translation initiation or initiation of mRNA degradation by RNaseE. We measured the effects of MicC overproduction on the *Salmonella hilD*’-’*lacZ* fusion in an *rne131* background strain, which has a defect in RNA degradosome assembly (8, 33-35). Although absolute expression of the *hilD* fusion was increased in the *rne131* background, MicC was still able to regulate (Fig. 5B). Therefore, we conclude that MicC base-pairing to the *hilD* mRNA blocks translation initiation but does not induce mRNA degradation.

### MicC expression is activated by SlyA and repressed by OmpR/EnvZ

Studies in *E. coli* (19) and transcriptomic analysis in *Salmonella* (21) suggested that MicC expression is repressed by OmpR/EnvZ and activated by SlyA. To characterize this regulation in more detail, we examined expression of a *micC’*-*lacZ*^*+*^ fusion. Deletion of either *envZ* or *ompR* in the fusion strain resulted in increased transcription of MicC in the exponential growth phase (Fig. 6A), confirming that the OmpR/EnvZ two-component system negatively regulates MicC. Deletion of *slyA*, in contrast, caused a 3-fold decrease in expression, showing that SlyA is a positive regulator of *micC* expression. Importantly, in the absence of SlyA, deletion of *ompR* or *envZ* had no effect, suggesting that OmpR/EnvZ function through, or are at least dependent on, SlyA for their control of *micC* transcription (Fig. 6A).

**FIG 6.**
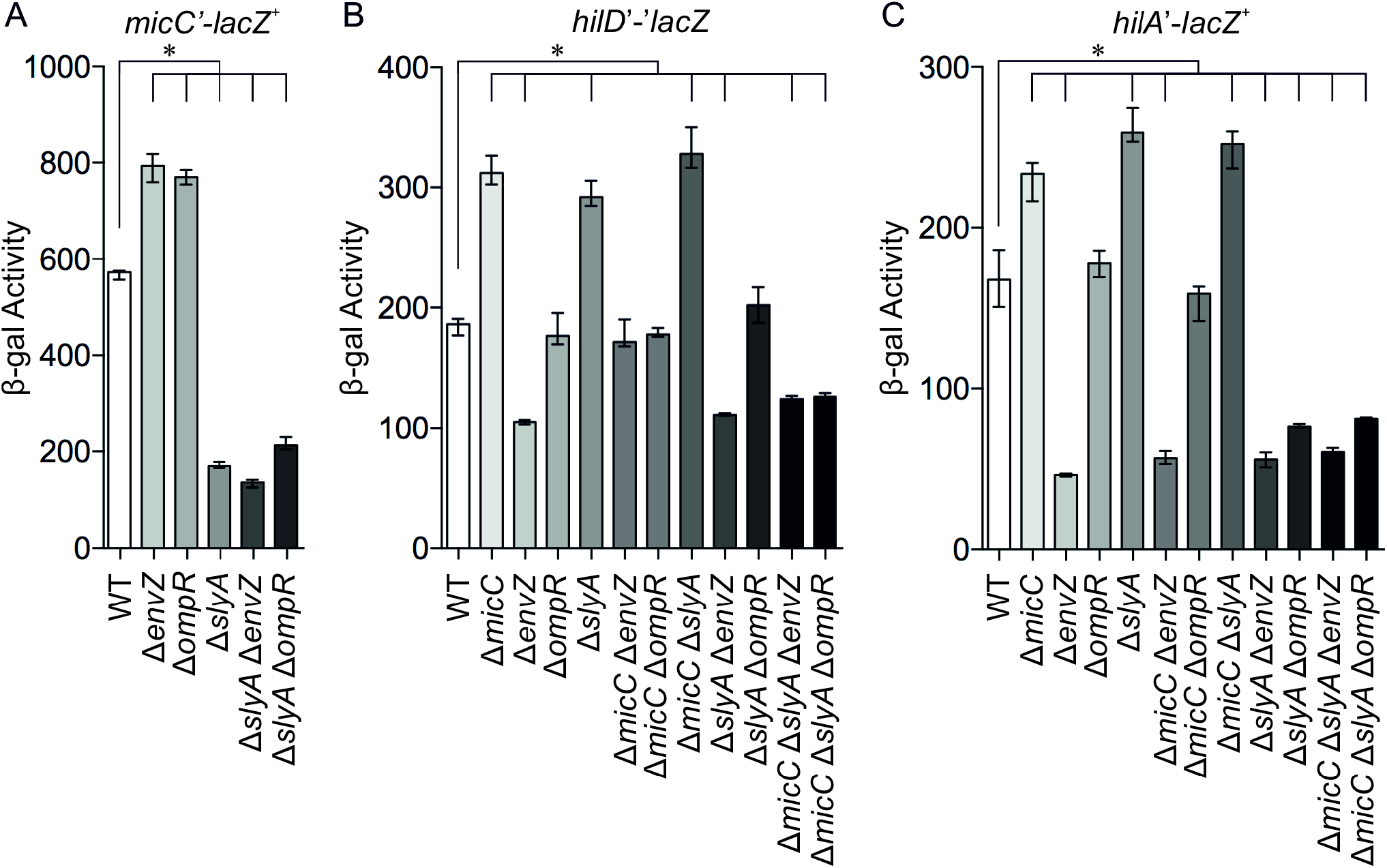
SlyA and EnvZ/OmpR regulate *micC* and the SPI1 T3SS. β-gal activity in *Salmonella* strains with a (A) *micC*’-*lacZ*^*+*^ transcriptional fusion, (B) *hilD*’-’*lacZ* translational fusion, or (C) *hilA*’-*lacZ*^*+*^ transcriptional fusion in backgrounds containing the indicated mutations. Results are shown as median with interquartile range and asterisks indicate significant differences between the datasets (n ≥ 4, *P* < 0.05, using a Kruskal–Wallis test followed by post hoc Dunn’s multiple comparisons). Strains used: JS749, JS892, JS2523, JS2524, JS2525, JS2526, JS2527, JS2528, JS2529, JS2530, JS2531, JS2532, JS2533, JS2534, JS2535, JS2536, JS2537, JS2538, JS2539, JS2540, JS2541, JS2542, JS2543, JS2544, JS2545, JS2546, JS2547, JS2548, JS2549 and JS2550.

To determine how this regulation affects SPI1 gene expression, we examined both a *hilD’*-‘*lacZ* translational (Fig. 6B) and *hilA*’-*lacZ*^+^ transcriptional fusions (Fig. 6C). Deletion of *micC* caused increased expression of both the *hilD* and *hilA* fusions, showing that MicC is affecting *hilD* translation under these conditions. As expected, deletion of *slyA* also led to an increase in expression of the *hilD* and *hilA* fusions, and deletion of *micC* in the *slyA* background had no further effect. These results show that SlyA is affecting SPI1 expression through MicC-mediated control of hilD translation.

Regulation by OmpR/EnvZ is more complicated. Deletion of *envZ* led to decreased expression of the *hilD’*-’*lacZ* fusion, but this decreased expression was also seen in the *envZ slyA micC* background (Fig 6B). It is interesting to note that deletion of *envZ* has a greater effect than mutations in *ompR*, as noted previously (7, 13). Note also that our Δ*ompR*::Cm allele is polar on the translationally coupled *envZ* (36). Thus, this decreased expression of the *hilD’*-’*lacZ* fusion seen in the *envZ* mutant is functioning through OmpR, but is independent of MicC.

Deletion of *envZ* also caused decreased expression of *hilA* (Fig 6C) that is independent of SlyA and MicC. Loss of OmpR (and EnvZ) has no effect under these conditions. Our previous data (7) showed that the primary effect of the *envZ* mutation in late stationary phase was to decrease HilD protein activity, leading to decreased expression of *hilA*. In the exponential phase data shown here, we also see an apparent effect on *hilD* transcription in the *envZ* mutant; we do not understand the mechanism. Thus OmpR/EnvZ, although controlling *micC* expression, also affect SPI1 independent of MicC, apparently through several mechanisms, which complicates interpretation of the data.

Transcriptomic data also implicated the sigma factor RpoS in the regulation of *micC* (21). As shown in Fig. S1A, deletion of *rpoS* caused increased expression of *micC* in early stationary phase but has no effect in exponential phase. Given that RpoS is acting negatively to control *micC*, this is likely an indirect effect. Deletion of *rpoS* also led to increased *hilD* transcription, particularly in stationary phase. This regulation was unaffected by loss of MicC (Fig. S1B). Thus, RpoS negatively regulates *hilD* by an unknown, and likely indirect mechanism (Fig. S1B), but this regulation is independent of MicC.

MicC is negatively regulated by OmpR/EnvZ and negatively regulates OmpC translation. Therefore, it was proposed to play a role in the differential osmoregulation of the porin proteins (19). As such, one would predict that MicC would be preferentially expressed at low osmolarity. We tested this hypothesis by examining expression of the *micC*’-*lacZ*^+^ fusion in low and high salt LB (Fig. S2). As shown in Fig. S2B, expression of *micC* is increased in high salt. Moreover, this regulation is largely independent of OmpR/EnvZ. Thus, overall regulation of *micC* is inconsistent with a simple role in regulation of the porins in response to osmolarity.

### Deletion of MicC enhances SPI1 dependent fitness in vivo

MicC regulates expression of the SPI1 T3SS via direct base pairing with the *hilD* mRNA. In vitro, this regulation is evident at mid-to late-exponential phase (Fig. 6). To determine if MicC affects SPI1 regulation in vivo in a manner that affects virulence, we performed competition assays using BALB/C mice. In oral infections, the Δ*micC* strain outcompeted the wildtype strain in both the upper small intestine (includes duodenum and jejunum) and lower small intestine (includes ileum) (Fig. 7A), the primary site of *Salmonella* invasion into epithelium cells (2, 3, 37). There was no significant fitness advantage for bacteria recovered from the spleen after either oral or intraperitoneal (IP) infection (Fig. 7B); systemic infection does not require SPI1 (5, 38). To determine whether the observed effects in the intestine were due to changes in SPI1 expression, we also performed competition assays in strains lacking the SPI1 T3SS. In the Δ*spi1* background, deletion of *micC* had no significant effect in the competition assay (Fig. 7C). These data are consistent with MicC having a significant regulatory role on *hilD* translation during intestinal infection.

**FIG 7.**
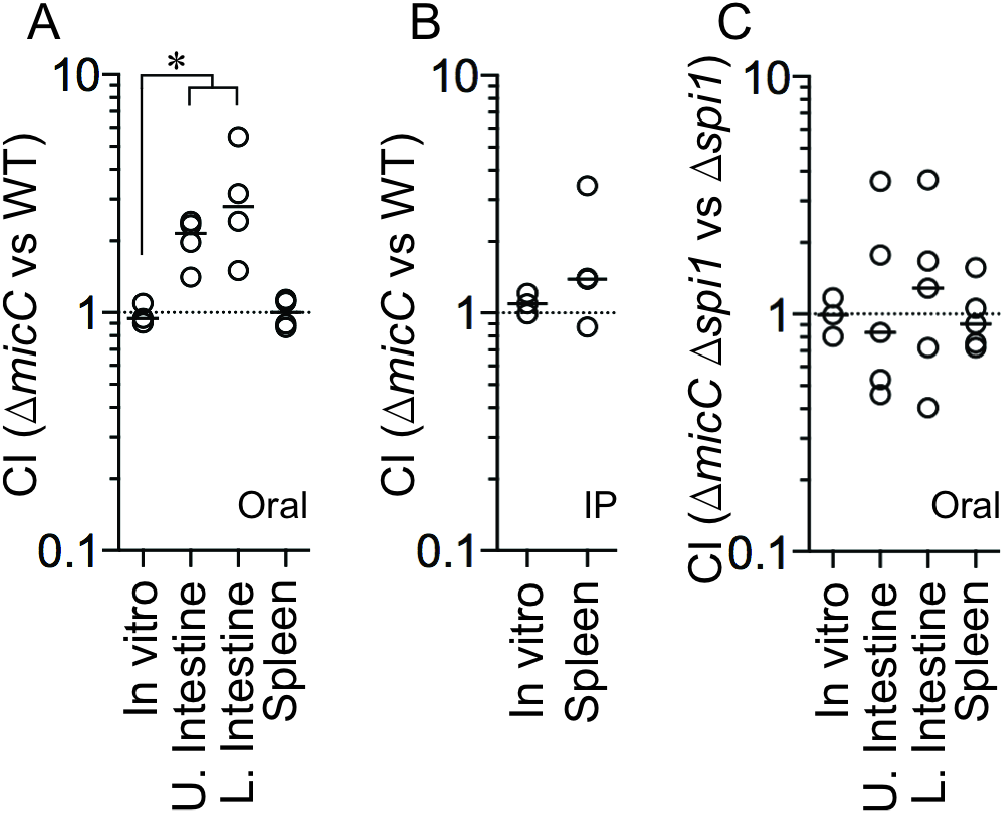
Deletion of MicC provides a fitness advantage in vivo. Competitive index (CI) for in vitro and in vivo infections comparing the following strains: (A) Δ*micC* to WT after oral infection, (B) Δ*micC* to WT after intraperitoneal (IP) infection or (C) Δ*micC* Δ*spi1* to Δ*spi1* after oral infection. Upper small intestine (contains duodenum and jejunum), lower small intestine (contains ileum) and spleen were harvested after oral infections, whereas only the spleen was harvested after IP infections. Each circle represents the CI from a single mouse. For in vitro competitions, N=3; panel A, N=4; panels B and C, N=5. The horizontal bars indicate the median of each dataset, and the asterisk indicates significant differences (*P* < 0.05) using a Mann-Whitney test. Strains used: JS135, JS2553, JS2554 and JS2555.

## DISCUSSION

The SPI1 T3SS is regulated in response to a plethora of physiological and environmental factors to ensure that this critical virulence factor is expressed at the appropriate time and place in the host and to optimize that expression. In this study, we identified the sRNA, MicC, as a repressor of *hilD* translation. MicC was previously identified as a negative regulator of the outer membrane porin proteins OmpC (19) and OmpD (20). MicC acts in the canonical fashion to control *hilD* translation, base pairing just upstream of the ribosome binding site (Fig. 3A). This binding blocks translation per se rather than initiating mRNA degradation (Fig. 3B). MicC also base pairs just upstream of the ribosome binding site in the *ompC* mRNA to block translation (19). Interestingly, in the case of *ompD*, MicC base pairs starting at 67 nucleotides downstream of the AUG and acts by initiating mRNA degradation versus blocking translation (20). MicC does not directly regulate *hilC, rtsA* or *hilA* (Fig. 4), showing that MicC-mediated downregulation of SPI1 T3SS is solely through regulation of *hilD* mRNA translation. These data reinforce HilD as the primary site of signal integration in the SPI1 regulatory circuit (Fig. 1).

Expression data in *E. coli* and transcriptome data in *Salmonella* suggested that *micC* transcription is controlled by SlyA, OmpR/EnvZ, and RpoS (21). Our data suggest that the primary transcriptional activator of *micC* is SlyA (Fig. 6A). OmpR/EnvZ negatively regulate *micC* transcription by affecting SlyA activation. Whether this regulation is all occurring directly at the *micC* promoter will require further investigation, but SlyA often works in conjunction with other transcriptional regulators in the control of gene expression (23, 24, 27, 28). Comparison of the sequence upstream of *micC* in various Enterobacteriaceae reveals a strikingly conserved sequence between -31 and -50 from the transcription start site (Fig. 2A). This suggests that this sequence is important for regulation, but it matches neither the SlyA (39-41) nor the OmpR consensus sequence (42). We also show that RpoS negatively regulates *micC* transcription in early stationary phase. This is almost certainly an indirect effect and determining the mechanism will also require further analyses.

The physiological signals that influence SlyA are unclear, although salicylate binds to and inactivates SlyA, and, and loss of SlyA affects the overall response to reactive oxygen species (25, 43, 44). SlyA is strongly induced when *Salmonella* is replicating in macrophages and *slyA* mutants are not able to survive in macrophages and are, therefore, attenuated for virulence (28). Our data show that SlyA increases the expression of MicC, which helps to repress the SPI1 T3SS. This regulation is apparently evident in the intestine with the *micC* mutant outcompeting the wildtype, consistent with increased expression of the SPI1 T3SS. The SPI1 T3SS is neither expressed nor required during systemic infection and replication in macrophages (5, 45). Our data suggest that this strong negative regulation of the system is mediated primarily by PhoPQ (46), but activation of MicC by SlyA may contribute to the downregulation of *hilD* transcription in the intracellular environment.

MicC, which is negatively regulated by OmpR and blocks *ompC* translation, was proposed to facilitate regulation of OmpF and OmpC in response to osmolarity (19). OmpR/EnvZ differentially regulate transcription of the porin genes such that *ompF* is preferentially transcribed in low osmolarity and *ompC* is preferentially transcribed in high osmolarity (12, 47). The sRNA MicF is co-transcribed with *ompC* and blocks *ompF* translation (18). Logic would dictate that MicC should be produced preferentially in low osmolarity to down regulate OmpC expression under these conditions. Interestingly, our results show that *micC* transcription increases with osmolarity (at least under the conditions we examined; Fig. S2) and that this regulation is independent of OmpR. Thus, the simple model does not hold. Indeed, OmpR is now known to be a global transcriptional regulator and most genes in the OmpR regulon are not osmoregulated (48), but rather activated at some threshold level of OmpR-P. Only if that threshold level is high, as in the case of *ompC*, is the gene preferentially activated at high osmolarity. Transcriptional regulation of *ompF* is more complex and apparently unique, being activated at low levels of OmpR-P, but then actively repressed by OmpR-P at higher levels (47-49). Negative regulation of *micC* by OmpR/EnvZ is via SlyA and the overall osmoregulation of *micC* is independent of the two-component system.

OmpR/EnvZ regulation of SPI1 is more complicated and one can argue that low levels of OmpR-P play a role. Deletion of EnvZ leads to a significant decrease in *hilA* transcription (Fig. 6). This effect functions through OmpR; the *ompR* mutation is polar on *envZ* (36). It has long been known that loss of EnvZ, but not OmpR/EnvZ, affects SPI1 expression (5, 7, 14). We previously showed that this *envZ* phenotype is mediated through control of HilD protein activity (7), which is consistent with the data shown here. However, those previous experiments were performed in late stationary phase. In the exponential phase experiments shown here, we also see an effect on *hilD* expression in the *hilD* null strain. In *E. coli*, there are low levels of OmpR-P in the *envZ* null strain, due to phosphorylation of OmpR by acetyl phosphate (50-52). Thus, it appears that low levels of OmpR-P are actively leading to decreased HilD protein activity (7) and perhaps *hilD* transcription/translation through unknown mechanisms. This is consistent with overall activation of SPI1 in high osmolarity, which would further be enhanced by OmpR-mediated repression of *micC*.

These results emphasize the role of HilD as the major signal integration point for control of the SPI1 T3SS. Most regulatory input is via post-transcriptional control of HilD, affecting HilD activity via protein-protein interactions, *hilD* translation, or mRNA stability (6-8, 29, 53), although the mechanisms are understood in only a few cases. We know that *hilD* translation is affected by binding of the RNA binding protein CsrA in the 39 nt *hilD* 5’ UTR (29, 54). Translation initiation is also controlled by the sRNAs FnrS, ArcZ (8) and MicC, all of which base pair at the ribosome binding site. All three of these sRNAs affect SPI1 expression during intestinal infection in the animal. More detailed analyses are required to determine the mechanisms and physiological role of additional sRNAs identified as affecting overall control of the T3SS (8).

## MATERIAL AND METHODS

### Bacterial strains and growth conditions

Bacterial strains and plasmids used in this study are described in Table S1. All *Salmonella* enterica serovar Typhimurium strains are isogenic derivatives of strain 14028 [American Type Culture Collection (ATCC)] and were constructed using P22 HT105/1 int-201 (P22)-mediated transduction. Deletion of genes or insertion of antibiotic resistance cassettes was performed using λ-red mediated recombination (55, 56). Insertions and deletions were confirmed by PCR and mutations were transferred into unmutagenized backgrounds by P22 transduction. In some cases, antibiotic resistance cassettes were removed using the temperature-sensitive plasmid pCP20 carrying the FLP recombinase (57). To create transcriptional *lacZ* fusions to MicC, the insertion mutation in MicC was converted to a transcriptional *lac* fusion using FLP-mediated recombination with plasmid pKG136, as previously described (56). The translational *lacZ* reporter fusions in *E. coli* were constructed using lambda Red-mediated recombination in strain PM1205, as described previously (8, 31).

Strains were routinely cultured in lysogeny broth (LB; 10% tryptone, 5% yeast extract, 5% NaCl). For SPI1 expression experiments, cells were grown in No Salt LB (NSLB; 10% tryptone, 5% yeast extract, 0% NaCl) or High Salt LB (HSLB; 10% tryptone, 5% yeast extract, 10% NaCl). All strains were grown at 37°C with aeration, except for the strains containing the temperature-sensitive plasmid pCP20 or pKD46, which were grown at 30°C. Antibiotics were used at the following final concentrations: ampicillin (Ap, 100 μg/mL), kanamycin (Km, 50 μg/mL), chloramphenicol (Cm, 20 μg/mL), apramycin (Apr, 50 μg/mL) and tetracycline (Tet, 10 μg/mL).

### Construction of plasmids and site-directed mutagenesis

The MicC small RNA was amplified from the *S*. Typhimurium 14028 genome using oligonucleotides pairs F-AatII-MicC and R-EcoRI-MicC and cloned into the pBR-p*lac* vector (31) after digestion with AatII and EcoRI restriction enzymes, creating pMicC. The *hilD* mRNA/MicC sRNA interactions were predicted using the IntaRNA RNA-RNA interaction tool (58). Mutations were introduced into the pMicC plasmid using a QuikChange Lightning site-directed mutagenesis kit (Stratagene). Oligonucleotides used in this study are listed in Table S2.

### ß-Galactosidase assays

ß-Galactosidase assays were performed using a microtiter plate assay as previously described (49). Briefly, *Salmonella* strains were inoculated in NSLB medium and grown ON at 37°C on a roller drum. These cultures were subsequently diluted 1:100 into 2 ml of HSLB medium and grown at 37°C on a roller drum for 3 hr or 8 hr (where indicated). For *E. coli* cultures, strains were initially inoculated into LB and grown overnight, then subcultured 1:100 into 2 ml of LB medium with 100 μM IPTG and 0.001% arabinose and grown at 37°C on a roller drum for 3 hr. ß-Galactosidase activity units are defined as [μmol of orthonitrophenol (ONP) formed min^-1^]×10^6^/(OD_600_ x ml of cell suspension).

### In vitro and in vivo competition assays

All animal work was reviewed and approved by the University of Illinois Institutional Animal Care and Use Committee (IACUC). Procedures were performed in our AAALAC accredited facility in accordance with University and PHS guidelines under protocol 15214. Competition assays in vivo and in vitro were performed using isogenic wild type and Δ*micC*, or ΔSPI1 and Δ*micC* ΔSPI1 mutants. Briefly, strains were grown overnight in LB. For oral infections, strains were mixed 1:1, washed, and suspended in 0.1 M phosphate buffered saline (pH 8) to an adjusted cfu ml^-1^ of 5×10^8^ (for wild type vs Δ*micC*) or 10^9^ (for ΔSPI1 vs Δ*micC* ΔSPI1). Before infection, food and water were withheld for 4 h, and then mice were inoculated with 200 μl of cell suspension by oral gavage. For intraperitoneal infections, 1:1 cell mixtures were diluted to 10^3^ cfu ml^-1^ in phosphate buffered saline (pH 7). Mice were inoculated with 200 μl of cell suspension by intraperitoneal (IP) injection. All inocula were diluted and plated on LB and then replica plated to appropriate antibiotic medium to determine the exact ratios of strains. After 3.5 days of infection, mice were sacrificed by CO_2_ asphyxiation and the spleens and small intestines were dissected from orally infected mice, or the spleens were dissected from IP infected mice. Tissues were mechanically homogenized in PBS with 15% glycerol and appropriate dilutions were plated on LB containing the appropriate antibiotics and subsequently replica plated to determine the ratio of strains recovered. In vitro competitions were performed simultaneously by subculturing 10^3^ cfu of the same inocula used for the in vivo experiments into 2 mL of LB. The cultures were incubated overnight at 37 °C with aeration, diluted, and plated on LB. Resulting colonies were replica plated onto LB containing the appropriate antibiotics. Results are presented as competitive index (CI), calculated as (percentage of strain A recovered/percentage of strain B recovered)/(percentage of strain A inoculated/percentage of strain B inoculated).

## ACKNOWLEDGEMENTS

This work was funded by NIH grant R01 GM120182 to JMS and CKV.

## Notes

### Competing Interest Statement

The authors have declared no competing interest.

