## Supplemental Material for "The sRNA MicC downregulates *hilD* translation to control the SPI1 T3SS in *Salmonella enterica* serovar Typhimurium"

#Corresponding Author

601 S Goodwin Avenue

Urbana Illinois, 61801

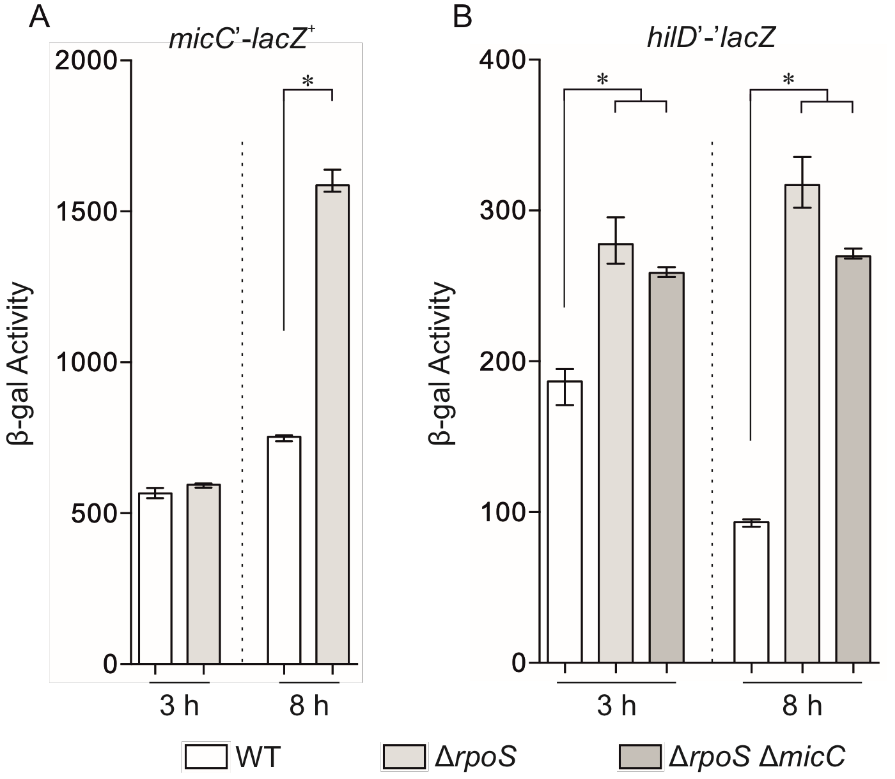

**FIG S1.** RpoS regulates MicC in stationary phase and the regulation of *hilD* by RpoS is independent of MicC. β-gal activity in *Salmonella* strains with a (A) *micC*'-*lacZ*$^+$ transcriptional fusion, or (B) *hilD*'-'*lacZ* translational fusion backgrounds containing the indicated mutations. Cells were grown in SPI1 inducing conditions for 3h (exponential phase) or 8h (stationary phase). Results are shown as median with interquartile range and asterisks indicate significant differences between the datasets (n = 4, $P < 0.05$, using a Mann-Whitney test). Strains used: JS892, JS2523, JS2556, JS2557 and JS2558.

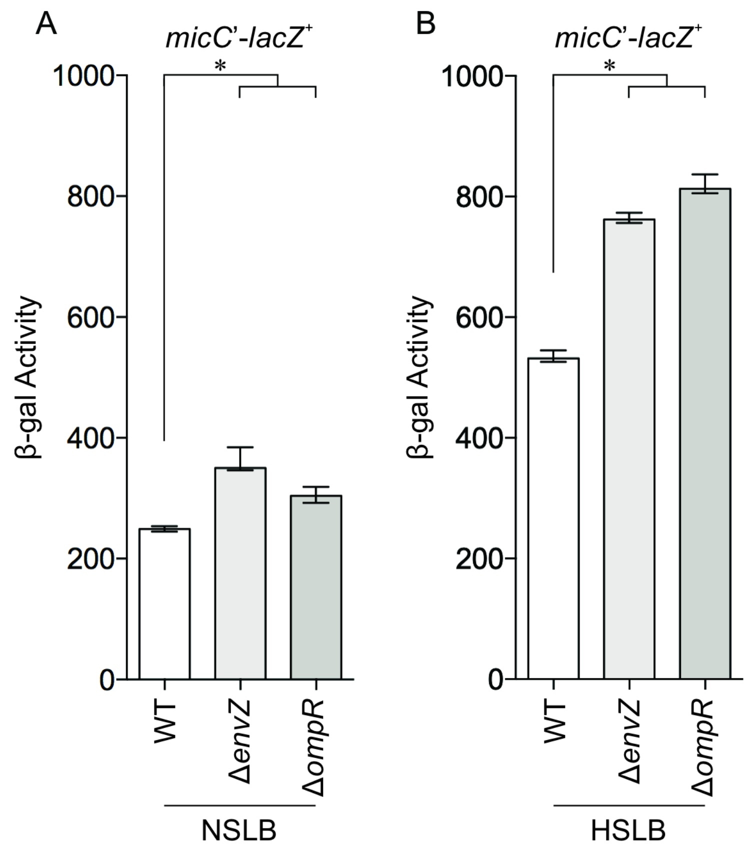

**FIG S2.** MicC expression is affected by salt independent of EnvZ/OmpR. β-gal activity in the *Salmonella micC*'-*lacZ*$^{+}$ transcriptional fusion strain with the indicated mutations grown in (A) no salt LB and (B) high salt LB. Results are shown as median with interquartile range and asterisks indicate significant differences between the datasets (n = 4, $P < 0.05$, using a Mann-Whitney test). Strains used: JS2523, JS2524 and JS2525.

**Table S1. Strains and plasmids used in this study**

| Strain or plasmid | Genotype | Deletion Endpoints[a] | Reference[b] |
|---|---|---|---|
| ***Salmonella* strains** | | | |
| 14028 | Wild type | | ATCC[c] |
| JS892 | Φ(*hilD*'-'*lacZ*)*hyb139* | | (1) |
| JS749 | *attλ*::*pDX1*::*hilA*'-*lacZ*$^+$ | | (2) |
| JS570 | Δ*hfq11*::Cm | | (3) |
| JS2118 | Δ*hfq11*::Cm Φ(*hilD*'-'*lacZ*)*hyb139* | | (4) |
| JS2117 | *rne131*::Cm | | (4) |
| JS2119 | *rne131*::Cm Φ(*hilD*'-'*lacZ*)*hyb139* | | (4) |
| JS2518 | Δ*micC*::Cm | 1,755,655-1,755,746 | |
| JS2519 | Δ*envZ*::Cm | 3,672,155-3,673,511 | |
| JS2520 | Δ*envZ*::Kan | 3,672,155-3,673,511 | |
| JS2521 | Δ*ompR*::Cm | 3,673,513-3,674,224 | |
| JS2522 | Δ*ompR*::Kan | 3,673,513-3,674,224 | |
| JS2180 | Δ*slyA*::Tet | | (5) |
| JS2523 | Φ(*micC*'-*lacZ*$^+$ ) | | |
| JS2524 | Δ*envZ*::Cm Φ(*micC*'-*lacZ*$^+$) | | |
| JS2525 | Δ*ompR*::Cm Φ(*micC*'-*lacZ*$^+$) | | |
| JS2526 | Δ*slyA*::Tet Φ(*micC*'-*lacZ*$^+$) | | |
| JS2527 | Δ*slyA*::Tet Δ*envZ*::Cm Φ(*micC*'-*lacZ*$^+$) | | |
| JS2528 | Δ*slyA*::Tet Δ*ompR*::Cm Φ(*micC*'-*lacZ*$^+$) | | |
| JS2529 | Δ*micC*::Cm Φ(*hilD*'-'*lacZ*)*hyb139* | | |
| JS2530 | Δ*envZ*::Cm Φ(*hilD*'-'*lacZ*)*hyb139* | | |
| JS2531 | Δ*ompR*::Cm Φ(*hilD*'-'*lacZ*)*hyb139* | | |
| JS2532 | Δ*slyA*::Tet Φ(*hilD*'-'*lacZ*)*hyb139* | | |
| JS2533 | Δ*micC* Δ*envZ*::Cm Φ(*hilD*'-'*lacZ*)*hyb139* | | |
| JS2534 | Δ*micC* Δ*ompR*::Cm Φ(*hilD*'-'*lacZ*)*hyb139* | | |
| JS2535 | Δ*micC* Δ*slyA*::Tet Φ(*hilD*'-'*lacZ*)*hyb139* | | |
| JS2536 | Δ*slyA*::Tet Δ*envZ*::Cm Φ(*hilD*'-'*lacZ*)*hyb139* | | |
| JS2537 | Δ*slyA*::Tet Δ*ompR*::Cm Φ(*hilD*'-'*lacZ*)*hyb139* | | |
| JS2538 | Δ*slyA*::Tet Δ*micC* Δ*envZ*::Cm Φ(*hilD*'-'*lacZ*)*hyb139* | | |
| JS2539 | Δ*slyA*::Tet Δ*micC* Δ*ompR*::Cm Φ(*hilD*'-'*lacZ*)*hyb139* | | |
| JS2540 | Δ*micC*::Cm *attλ*::*pDX1*::*hilA*'-*lacZ*$^+$ | | |
| JS2541 | Δ*envZ*::Cm *attλ*::*pDX1*::*hilA*'-*lacZ*$^+$ | | |
| JS2542 | Δ*ompR*::Cm *attλ*::*pDX1*::*hilA*'-*lacZ*$^+$ | | |
| JS2543 | Δ*slyA*::Tet *attλ*::*pDX1*::*hilA*'-*lacZ*$^+$ | | |
| JS2544 | Δ*micC*::Cm Δ*envZ*::Kan *attλ*::*pDX1*::*hilA*'-*lacZ*$^+$ | | |
| JS2545 | Δ*micC*::Cm Δ*ompR*::Kan *attλ*::*pDX1*::*hilA*'-*lacZ*$^+$ | | |
| JS2546 | Δ*micC*::Cm Δ*slyA*::Tet *attλ*::*pDX1*::*hilA*'-*lacZ*$^+$ | | |
| JS2547 | Δ*slyA*::Tet Δ*envZ*::Cm *attλ*::*pDX1*::*hilA*'-*lacZ*$^+$ | | |

| | | | |
|---|---|---|---|
| JS2548 | Δ*slyA*::Tet Δ*ompR*::Cm *attλ*::*pDX1*::*hilA*'-*lacZ*$^{+}$ | | |
| JS2549 | Δ*slyA*::Tet Δ*micC* Δ*envZ*::Cm *attλ*::*pDX1*::*hilA*'-*lacZ*$^{+}$ | | |
| JS2550 | Δ*slyA*::Tet Δ*micC* Δ*ompR*::Cm *attλ*::*pDX1*::*hilA*'-*lacZ*$^{+}$ | | |
| JS2196 | φ(*hilC*'-*lacZ*$^{+}$)*113* | | (5) |
| JS2187 | φ(*rtsABCD10*-*lacZ*$^{+}$) | | (5) |
| JS253 | Δ*hilD*114::Cm | | (6) |
| JS2551 | Δ*hilD*114::Cm φ(*hilC*'-*lacZ*$^{+}$)*113* | | |
| JS2552 | Δ*hilD*114::Cm φ(*rtsABCD10*-*lacZ*$^{+}$) | | |
| JS135 | *zii-8104*::Tn*10*dTc | | (7) |
| JS2553 | Δ*micC*::Cm *zii-8104*::Tn*10*dTc | | |
| JS481 | Δ*spi1-2916* | | (8) |
| JS2554 | Δ*spi1-2916 zii-8104*::Tn*10*dTc | | |
| JS2555 | Δ*micC*::Cm Δ*spi1-2916 zii-8104*::Tn*10*dTc | | |
| JS539 | Δ*rpoS*::Tet | | (9) |
| JS2556 | Δ*rpoS*::Tet Φ(*micC*'-*lacZ*$^{+}$) | | |
| JS2557 | Δ*rpoS*::Tet Φ(*hilD*'-'*lacZ*)*hyb139* | | |
| JS2558 | Δ*rpoS*::Tet Δ*micC*::Cm Φ(*hilD*'-'*lacZ*)*hyb139* | | |
| ***E. coli* strains** | | | |
| PM1205 | MG1655 *mal*::*lacI*$^{q}$, Δ*araBAD araC*$^{+}$, *lacI*'::P$_{BAD}$-*cat*-*sacB*:*lacZ*, *miniλtet*$^{R}$ | | (10) |
| JMS6500 | PM1205 *lacI*'::P$_{BAD}$-*hilD*'-'*lacZ* | | (4) |
| JMS6510 | PM1205 *lacI*'::P$_{BAD}$-*hilD*mt'-'*lacZ* | | |
| JMS6503 | PM1205 *lacI*'::P$_{BAD}$-*hilC*'-'*lacZ* | | (4) |
| JMS6504 | PM1205 *lacI*'::P$_{BAD}$-*rtsA*'-'*lacZ* | | (4) |
| JMS6505 | PM1205 *lacI*'::P$_{BAD}$-*hilA*'-'*lacZ* | | (4) |
| **Plasmids** | | | |
| pBRplac | p*lac* promoter-based expression vector, Ap$^{R}$ | | (11) |
| pMicC | pBRplac with MicC sRNA, Ap$^{R}$ | | |
| pMicC-mt | pBRplac with mutated MicC sRNA, Ap$^{R}$ | | |
| pKD3 | *bla* FRT *cat* FRT PS1 PS2 oriR6K, Ap$^{R}$ | | (12) |
| pKD4 | *bla* FRT *aph* FRT PS1 PS2 oriR6K, Ap$^{R}$ | | (12) |
| pKD46 | *bla* P$_{BAD}$*gam bet exo* PSC101 oriTS, Ap$^{R}$ | | (12) |
| pCP20 | *bla cat c*I857 λP$_{R}$*flp* pSC101 oriTS, Ap$^{R}$ | | (13) |
| pKG136 | *ahp* FRT *lacZY*$^{+}$ t$_{his}$ oriR6K, Ap$^{R}$ | | (14) |

a. Base pairs, inclusive, of deletions as in *S. enterica* serovar *Typhimurium* 14028 genome sequence (NCBI Genbank CP001363.1).

b. This study, unless otherwise noted

c. ATCC, American Type Culture Collection

**Table S2. List of oligonucleotides**

| Name of Primer | Sequence (5’ to 3’) |
|---|---|
| F-micC | TCGGTTACGATATTCTACGCCTGTTATATGCCTTTATTGTTGTAGGCTGGAGCTGCTTCG |
| R-micC | GGCGCAGATTAAAAAATATTCTAAGGATTAACCTGGAAACCATATGAATATCCTCCTTAG |
| F-envZ | CTACGTCTTTGTACCGGACGGTTCTAAAGCATGAGGCGATGTAGGCTGGAGCTGCTTCG |
| R-envZ | TCCGGCGTTGAGAAGAAAGGGAGGGTAATACCTCCCTTTCCATATGAATATCCTCCTTAG |
| F-ompR | CACACTTACATTTGTTGCGAACCTTTGGGAGTACAGACATGTAGGCTGGAGCTGCTTCG |
| R-ompR | TGAACTTCGCGGTGAGAAGCGCATTCGCCTCATGCTTTCATATGAATATCCTCCTTAG |
| F-micC-mt | AAGATACTGACGTCCAATAATGCCAAATTTGTCACATATTCA |
| R-micC-mt | TGAATATGTGACAAATTTGGCATTATTGGACGTCAGTATCTT |
